# Alpha-synuclein supports interferon stimulated gene expression in neurons

**DOI:** 10.1101/2020.04.25.061762

**Authors:** Aaron R. Massey, Brendan Monogue, Yixi Chen, Kelsey Lesteberg, Michaela E. Johnson, Liza Bergkvist, Jennifer A. Steiner, Jiyan Ma, Ravi Mahalingam, B.K. Kleinschmidt-Demasters, Martha L Escobar Galvis, Patrik Brundin, Tilo Kunath, J. David Beckham

## Abstract

The protein alpha-synuclein (asyn) is predominantly expressed in neurons and is associated with neurodegenerative diseases like Parkinson’s disease (PD); yet, a functional role for asyn in neurons is not clearly established. We have previously shown that asyn expression is up-regulated following viral infection in neurons and is critical for host immune responses to RNA virus infections. Here, we investigate the mechanisms underlying asyn-dependent immune responses to RNA virus infection in the brain. Using asyn knock-out (KO) mice and human neuronal models, we show that asyn is required for expression of the full repertoire of interferon-stimulated genes (ISGs) in neurons following acute RNA virus infection. Furthermore, treatment of asyn KO human neurons with poly I:C or type I interferon also fail to induce expression of the full complement of ISGs suggesting that asyn plays an important role in modulating neuronal innate immune responses. In brain tissue, asyn-dependent ISG expression is independent of microglia activation and supports activation of infiltrating lymphocytes following viral challenge. We also show that virus infections lead to accumulation of phosphorylated S129 asyn in human and non-human primate neuronal tissues. In a model of pS129 asyn pathology, we found that infection with West Nile virus increases microglia activation but does not significantly alter pS129 asyn pathology in the mouse model. Taken together, our results establish asyn as a novel, neuron-specific modulator of innate immunity by a mechanism that promotes interferon-stimulated gene expression and links responses to virus infection with formation of phosphorylated S129-asyn in neuronal tissue.

## Introduction

Previous studies have shown that asyn is up-regulated in neurons following viral infection and that knockout of the asyn gene, *Snca*, in mice results in increased viral growth in the brain and increased mortality; yet, the mechanism for asyn-dependent protection from virus infection remains undefined (Bantle et al., 2019; Beatman et al., 2015; Stolzenberg et al., 2017). We previously reported that asyn knockout (KO) mice (*Snca* -/-) exhibit increased disease severity and viral growth in the brain following peripheral challenge with RNA viruses including West Nile virus (WNV) and Venezuelan equine encephalitis virus (VEEV) TC83(Beatman et al., 2015). An important subsequent study demonstrated that total asyn expression was elevated in gastrointestinal-associated neurons following viral gastroenteritis in children(Stolzenberg et al., 2017). The same study also suggested that asyn expression supported chemotaxis and activation of infiltrating dendritic cells(Stolzenberg et al., 2017). More recently, intranasal inoculation of Western equine encephalitis virus (WEEV) was shown to result in viral spread along olfactory neuronal pathways of mice, resulting in activation of microglia and astrocytes in multiple brain regions, including the midbrain and striatum(Bantle et al., 2019). WEEV infection also resulted in loss of dopaminergic neurons in the substantia nigra and formation of prominent proteinase K-resistant aggregates of phospho-serine129 modified asyn (Bantle et al., 2019). Taken together, recent studies have shown that acute RNA virus infection increases asyn levels in mice, increases phosphorylation of asyn at serine residue 129, and can result in loss of nigral dopamine neurons; thereby recapitulating multiple key neuropathological features of PD. Despite these important recent findings, links between asyn expression and a functional role in the immune responses of the central nervous system remain unclear. We now show that asyn mediates ISG expression in neurons downstream of interferon signaling resulting in activation of infiltrating of T-cells, and that viral infection increases levels of phosphorylated S129 asyn in brain tissue.

## Results

### Brain tissue from asyn KO mice exhibit diminished ISG responses following WNV challenge

Our previous work has showed that asyn KO mice exhibited increased mortality and increased viral growth in the central nervous system (CNS) compared to WT mice when challenged with WNV and VEEV TC83(Beatman et al., 2015). In WT mice, the presence of asyn expression restricted virus in neuronal tissue and viral growth in the spleen was not altered by the presence of asyn (Beatman et al., 2015). Based on these data, we evaluated the brains of asyn KO mice for evidence of decreased gene expression of immune-associated genes despite a higher virus burden compared to WT mice.

To define specific immune pathways in the CNS that may be involved in asyn-dependent viral immune responses, we inoculated asyn wild-type (WT, *Snca* +/+) and asyn KO (*Snca* -/-) mice with mock diluent or WNV (1000pfu, subcutaneous inoculation). At day 8 post-infection, we harvested brain tissue and analyzed total RNA using RNAseq. Time points past day 8 are not possible since asyn KO mice exhibit high mortality rates following this day as previously described(Beatman et al., 2015). Mock-inoculated WT and asyn KO mice exhibited similar gene expression profiles, and large sets of genes were upregulated in brain tissue following WNV infection of WT and asyn KO mice (**Fig. 1A**). Analysis of differentially regulated genes in WNV-infected, asyn KO brain tissue compared to WNV-infected WT brain tissue revealed a clear subset of 39 genes that exhibited significantly decreased expression in virus infected asyn KO brain tissue compared to WT brain tissue (**Fig. 1B, Table**). Many of the identified genes were known to be involved in virus infections and the immune response including Usp11, Tmprss2, Prrx1, and Hsp90b1(Hoffmann et al., 2020; Istomine et al., 2019; Jacko et al., 2016; Limburg et al., 2019; Wang et al., 2018). Upon further analysis of gene expression as measured by mean FPKM (Fragments per Kilobase of transcript per Million), we found that *Oas1b* expression was significantly decreased by 50% in WNV-infected asyn KO mouse brain tissue compared to WNV-infected WT mouse brain tissue (**Fig. 1C**). Similarly, we found evidence of a 50% decrease in mean FPKM in other ISGs important in the antiviral responses including *Ifit2* and *Trim25* (**Fig. 1D,F).**

**Fig. 1.**
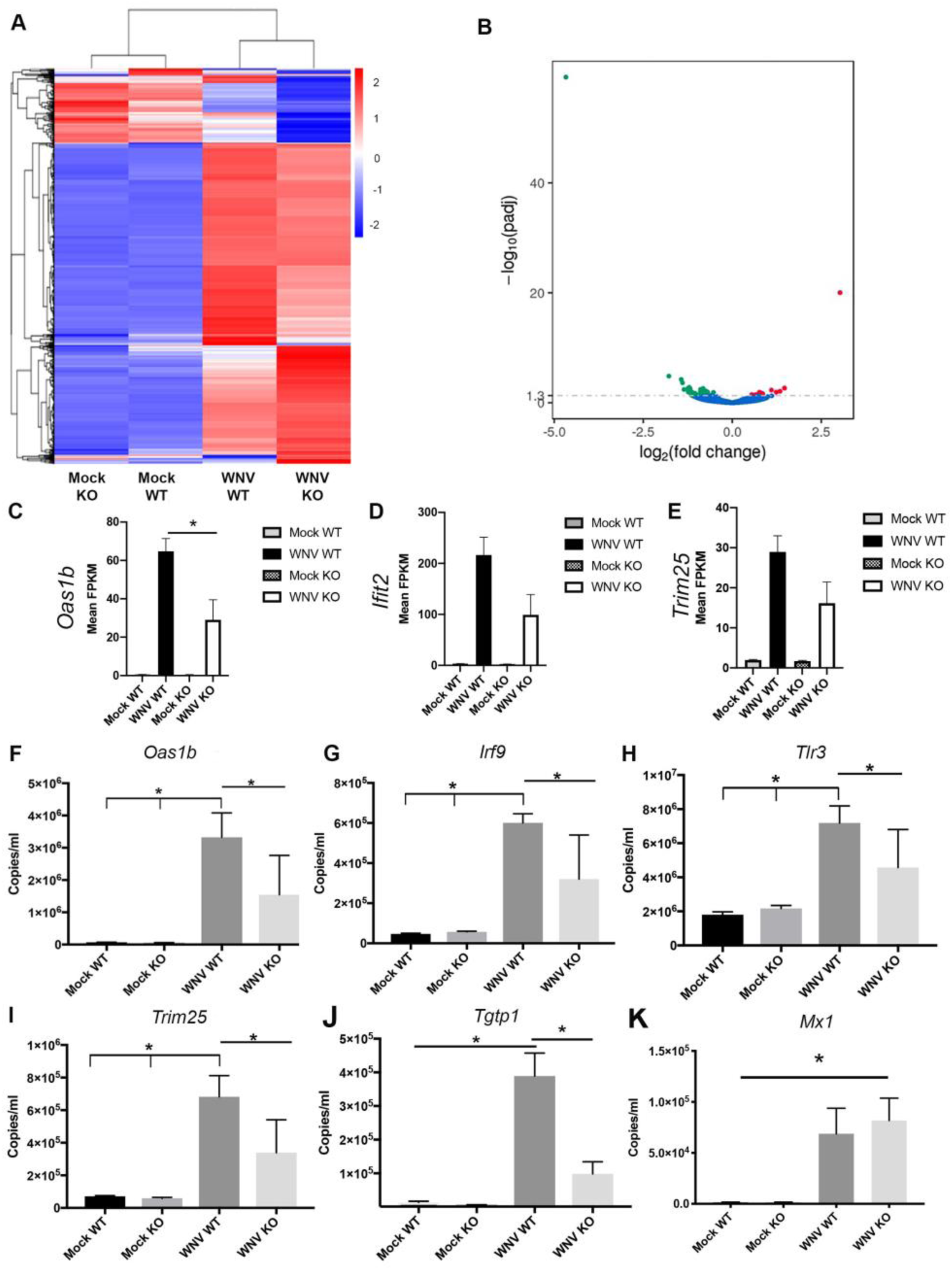
Alpha-synuclein knockout mice exhibit decreased expression of specific ISGs in the brain following WNV infection. **A**) RNAseq heat map cluster analysis of differential expression of genes in the brain from indicated treatment groups using hierarchical clusters of log10(FPKM+1). n=4 mice per group. **B**) Graphic representation of differential gene expression comparing WNV-infected WT and KO brain tissue. Green: genes with significantly decreased expression in KO brain tissue. Red: genes with significantly increased gene expression in KO brain tissue. Mean FPKM values for **C**) *Oas1b*, **D)** *Ifit2*, and **E)** *Trim25* are reduced in WNV-infected KO brain tissue compared to WT brain tissue. To validate RNAseq data, QPCR values from brain tissue is shown in indicated treatment groups to quantify expression of ISGs identified in RNAseq analysis including **F**) *Oas1b*, **G**) *Irf9*, **H**) *Tlr3*, **I**) *Trim25*, and **J**) *TGTP1*. **K**) QPCR values for *Mx1* in brains of mice as an example that only a subset of ISGs in the brain are regulated by asyn expression. n=3 mice per treatment group. WT=asyn wild-type mice, KO=asyn knockout mice. *p<0.01. ANOVA.

To validate our findings showing decreased ISG responses in the brains tissue of WNV-infected asyn KO mice despite robust viral replication, we performed RT-qPCR on brain tissue from the same treatment groups for specific ISGs involved in important antiviral responses. We found that gene expression of *Oas1b, Irf9, Tlr3, Trim25, and Tgtp1* exhibited significantly decreased expression in the brain tissue from WNV-infected, asyn KO mice compared to WNV-infected, WT mice (**Fig. 1F-J**). Notably, this was a subset of genes and there were many inflammatory pathway genes that were activated by WNV infection in both WT and KO mouse brain tissue including the antiviral gene *Mx1* (**Fig. 1K**). These data suggest that asyn KO mice are more susceptible to viral infection due to a deficient ISG response in the brain.

### Interferon Stimulated Gene (ISG) expression is dependent on alpha-synuclein in primary neurons

Next, we determined if neurons require asyn for ISG expression in a cell autonomous manner. Since RNA isolated from brain tissue represents a highly complex mixture of neuronal and non-neuronal cells, we utilized a highly pure population of human dopaminergic neurons differentiated from *SNCA* +/+ (WT) and *SNCA* -/- (KO) human embryonic stem cells (hESCs, **Fig. 2A)**(Chen et al., 2019). Since WNV infection results in rapid death of primary neurons in culture, we utilized VEEV TC83, an attenuated RNA virus that infects neurons. Also, TC83 virus is an attenuated virus due in part to a mutation in the 5’ untranslated region that disrupts a RNA stem-loop that is required to escape IFIT1 restriction(Reynaud et al., 2015). Thus, TC83 is able to infect and replicate at levels in neurons with decreased expression of IFIT1 and related innate immune responses.

**Fig. 2.**
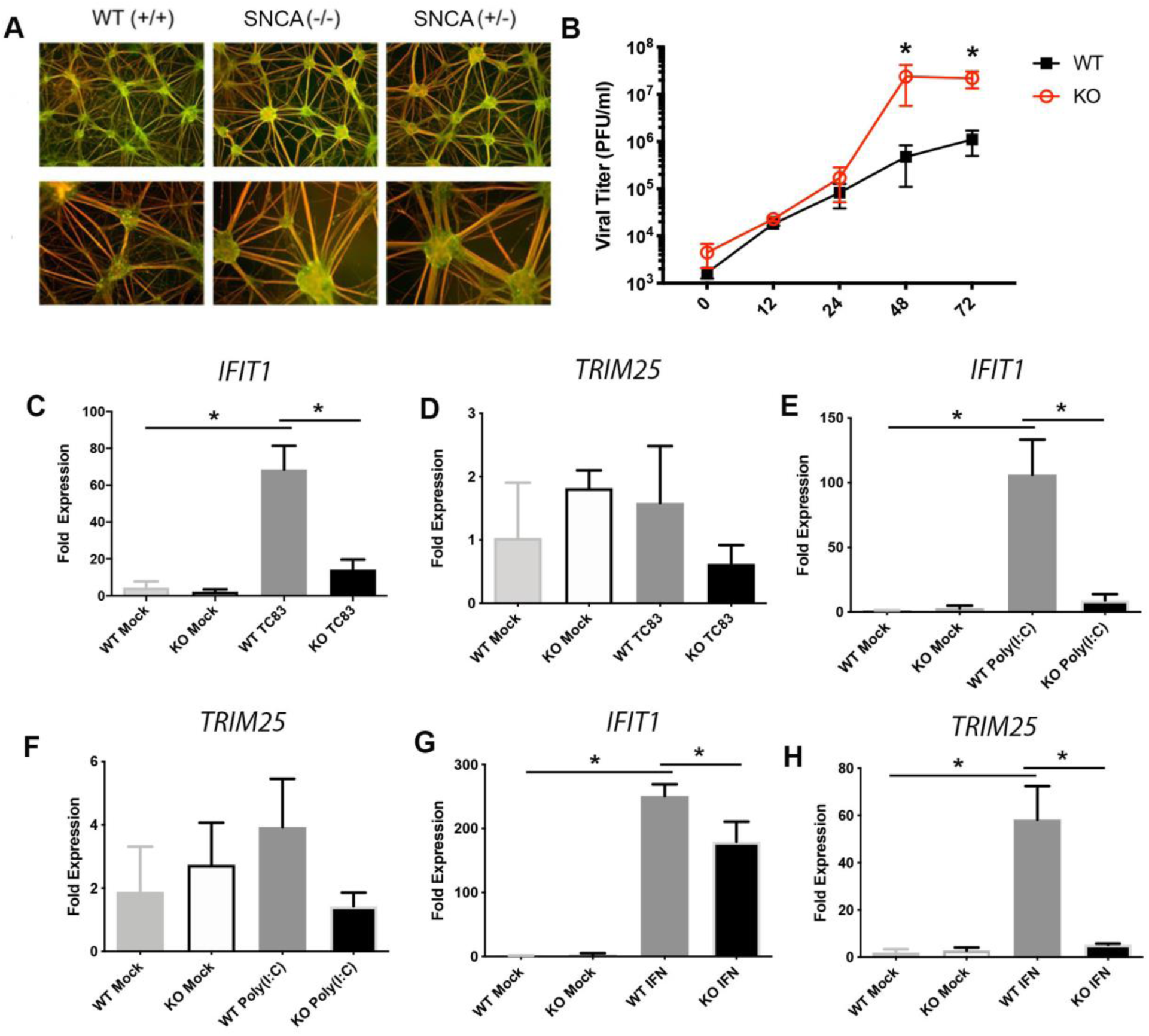
Alpha-synuclein expression supports interferon-stimulated gene expression in primary neurons. **A**) CRISPR/Cas-induced knockout of one or two *SNCA* genes in embryonic stem cells (ESC) was completed followed by differentiation into primary human neurons positive for beta-III tubulin (red) and tyrosine hydroxylase (green). Images shown at day 42 in culture. **B**) Asyn WT and knockout (KO) neurons were inoculated with VEEV TC83 (MOI 1) and supernatant collected at indicated time points for viral titer assay. *p<0.05, two-way ANOVA. N=4 replicates per group. Next, asyn WT and KO neurons were inoculated with mock or VEEV TC83 (MOI 10) and neurons were harvested at 12 hours post-infection for gene expression analysis of **C**) *IFIT1* and **D**)*TRIM25* expressed a fold-increase in gene expression compared to mock-inoculated WT neurons. *p<0.0001, one-way ANOVA, multiple comparisons. N=4-5 replicates per group. Asyn WT and KO neurons were treated with poly I:C (25 microg/ml) and neurons harvested at 4 hours post-treatment for **E**) *IFIT1* **F**) *TRIM25* gene expression analysis. N=3-4 per group. Asyn WT and KO neurons were treated with type 1 interferon (1000IU/ml) and cells harvested at 4 hours post-treatment for **G**) *IFIT1* and **H**) *TRIM25* gene expression analysis. *p=0.007, two-way ANOVA, multiple comparisons. N=3 per group.

Human asyn WT and asyn KO neurons were inoculated with TC83 (MOI 1) and supernatant collected over 72 hours. We found that TC83 viral titer was significantly increased in asyn KO neurons compared to WT neurons (**Fig. 2B**). Next, we determined if loss of asyn expression altered ISG expression in human neurons following virus infection. Human WT and asyn KO neurons were inoculated with TC83 (MOI 10), cells were harvested at 12 hours post-infection, and total RNA isolated. RT-qPCR analysis for gene expression for *IFIT1* and *TRIM25* expression revealed that TC83-infected, asyn KO neurons exhibit significantly decreased gene expression of *IFIT1* compared to TC83-infected asyn WT neurons despite similar viral loads at this early time point (**Fig. 2C**). Since *IFIT1* is an important restriction factor for VEEV TC83 and mutations in TC83 prevent *IFIT1* escape, then it follows that asyn KO neurons with decreased *IFIT1* expression would allow for increased viral growth. Interestingly, TC83 infection did not significantly induce *TRIM25* in this human neuronal system such that no difference in *TRIM25* expression was detected between virus-infected WT and asyn KO neurons (**Fig. 2D**).

Next, we determined if changes in ISG expression were related to RNA sensing mechanisms and type 1 interferon signaling. Human asyn KO and WT neurons were treated with control diluent, Poly I:C, or type 1 interferon, and neurons were harvested at 4 hours post-treatment for RNA analysis using QPCR for *IFIT1* and *TRIM25* expression. We found that asyn KO neurons exhibit significantly decreased gene expression of *IFIT1* following poly I:C treatment while *TRIM25* expression was not significantly induced in human neurons (**Fig. 2E,F**). Next, we found significantly decreased expression of both *IFIT1* and *TRIM25* in asyn KO neurons compared to WT neurons following type 1 interferon treatment (**Fig. 2G,H**). These data show for the first time that cell autonomous asyn expression supports ISG expression in neurons following RNA virus infection, RNA sensing & signaling, and type 1 interferon stimulation.

### Asyn expression does not alter virus-induced microglia activation in the brain

Our data show that neuronal expression of asyn modulates expression of neuronal interferon stimulated genes. Next, we determined the effect of asyn-dependent ISG expression on other cell types in the brain. Previous studies have shown that neuronal asyn expression can activate microglia and promote an inflammatory response(Harms et al., 2013; Roodveldt et al., 2013; Schapansky et al., 2015). Other studies indicate asyn-dependent activation of microglia occurs through a TLR-dependent mechanism (Beraud et al., 2013; Beraud et al., 2011; Caplan and Maguire-Zeiss, 2018; Roodveldt et al., 2010; Roodveldt et al., 2013; Sanchez-Guajardo et al., 2015). Using both WNV and TC83 virus infections, we first determined the role of asyn expression on the activation of microglia in the brain.

First, we inoculated asyn wild-type (WT, *Snca* +/+) and asyn KO (*Snca* -/-) mice with mock diluent or WNV (1000pfu, subcutaneous inoculation). At day 8 post-infection, we harvested brain tissue for histologic analysis for Iba1 and GFAP immunohistochemistry. We found that Iba1+ cells increased expression and number in both WNV-infected WT and asyn KO brain tissue (**Fig. 3A**). Using immunofluorescence for Iba1 and GFAP in the same tissue samples, we found that both microglia and astrocyte activation, respectively, in the brains of WNV-infected asyn KO and WT mice, exhibit no significant differences in activation between the two genotypes (**Fig. 3A-D**). These data show that WNV infection results in microglia and astrocyte activation independent of asyn expression.

**Fig. 3.**
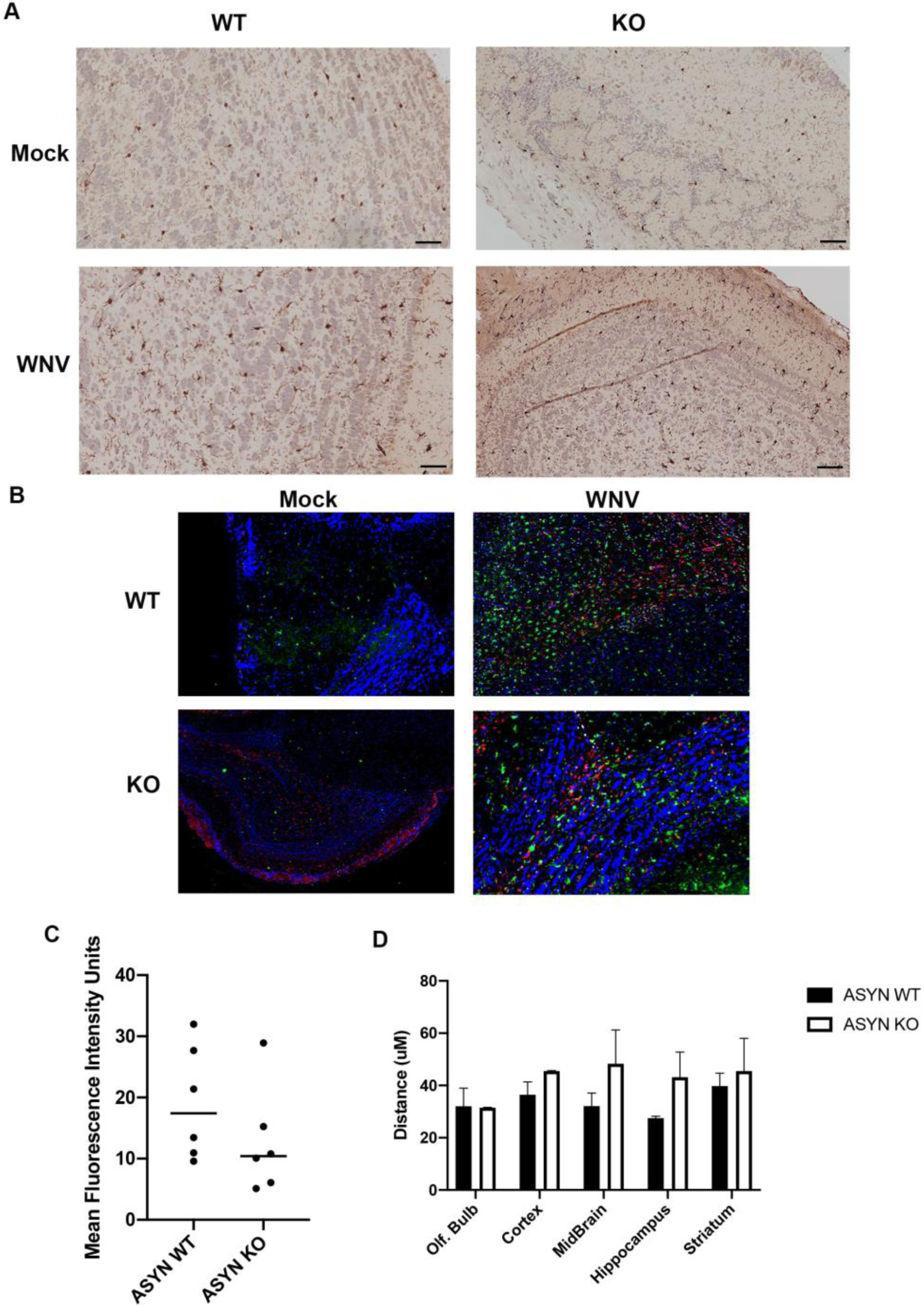
West Nile virus increases microglia activation independent of ASYN expression. ASYN WT and KO mice were inoculated with WNV (1000pfu, subcutaneous inoculation) and brain tissue harvested for analysis at day 8 post-infection. **A**) Representative histology images of the cortex labeled with antibody to Iba1 in the indicated treatment groups indicates increased Iba1 staining following WNV infection in both ASYN WT and KO treatment groups. N=3. **B**) Immunofluorescence imaging from the same treatment groups showing representative images from the midbrain showing nuclei (DAPI, blue), microglia (Iba1, green), astrocytes (GFAP, red), and WNV envelope antigen (white) showing evidence of increased expression of Iba1 and GFAP following WNV infection in both treatment groups. N=3 **C**) Quantification of mean fluorescence intensity (MFI) units per high power field of GFAP expression following WNV infection in the indicated treatment groups. p=0.24. Mean MFI obtained from N=6 mice per group. **D**) In each brain sample, two-point distance measured from WNV antigen-infected neurons to Iba1+ microglia and mean distance provided per brain section for WNV-infected ASYN WT and KO mice. N=3 mice per group. p=0.488, Mixed effects model, multiple comparisons.

Next, we evaluated the role of TC83 virus infection in activation of microglia in WT and KO mice. We have previously shown that TC83 infects the neurons of the CNS in asyn KO mice but not in WT mice(Beatman et al., 2015). To normalize for TC83 virus infection in the brain, we established a model to study acute central nervous system microglia activation following intracerebral inoculation of TC83 virus. We injected asyn WT and asyn KO mice by intracerebral inoculation with VEEV TC83 (TC83, 1000pfu). Following intracerebral inoculation, TC83-infected asyn KO mice exhibited significantly increased weight loss compared to TC83-infected asyn WT mice (**Fig. 4A**). At day 4 post-infection, mice were sacrificed and plaque assay analysis revealed significantly increased TC83 virus titer in the brains of asyn KO mice compared to WT mice (**Fig. 4B**). We next determined the role of asyn in activation of microglia in the brain following intracerebral injection of asyn WT and KO mice with TC83 (1000pfu, intracerebral injection). Brain tissue was harvested at day 4 post-infection as a time point with significant viral growth in the brain, but a time-point prior to significant peripheral inflammatory cell infiltration. Flow cytometry analysis of brain tissue revealed significantly increased numbers of CD45+CD11b+CD68+ cells, CD45+CD11b+IL6+ cells, and CD45+CD11b+TNFalpha+ following TC83 infection in both WT and asyn KO brain tissue, but we found no difference in CD45+CD11b+ cell activation when comparing asyn WT and KO brain tissue (**Fig. 4C-E, Fig. S1**). Next, we determined if microglial gene expression following TC83 infection in the brain was dependent on asyn expression. Using the same TC83 infection experimental model as above, microglia were column isolated from brain tissue using CD11b antibody binding. Total RNA was extracted from CD11b+ brain cells and analyzed by PCR array for ISG expression (**Fig. 4F**). ISG expression was normalized to mock-infected, WT microglia and revealed similar increases in ISGs following VEEV TC83 infection in both WT and asyn KO mice. Based on these data, we conclude that asyn-dependent regulation of innate immune responses and ISG responses were not due to differences in microglia activation following virus infection.

**Fig. 4.**
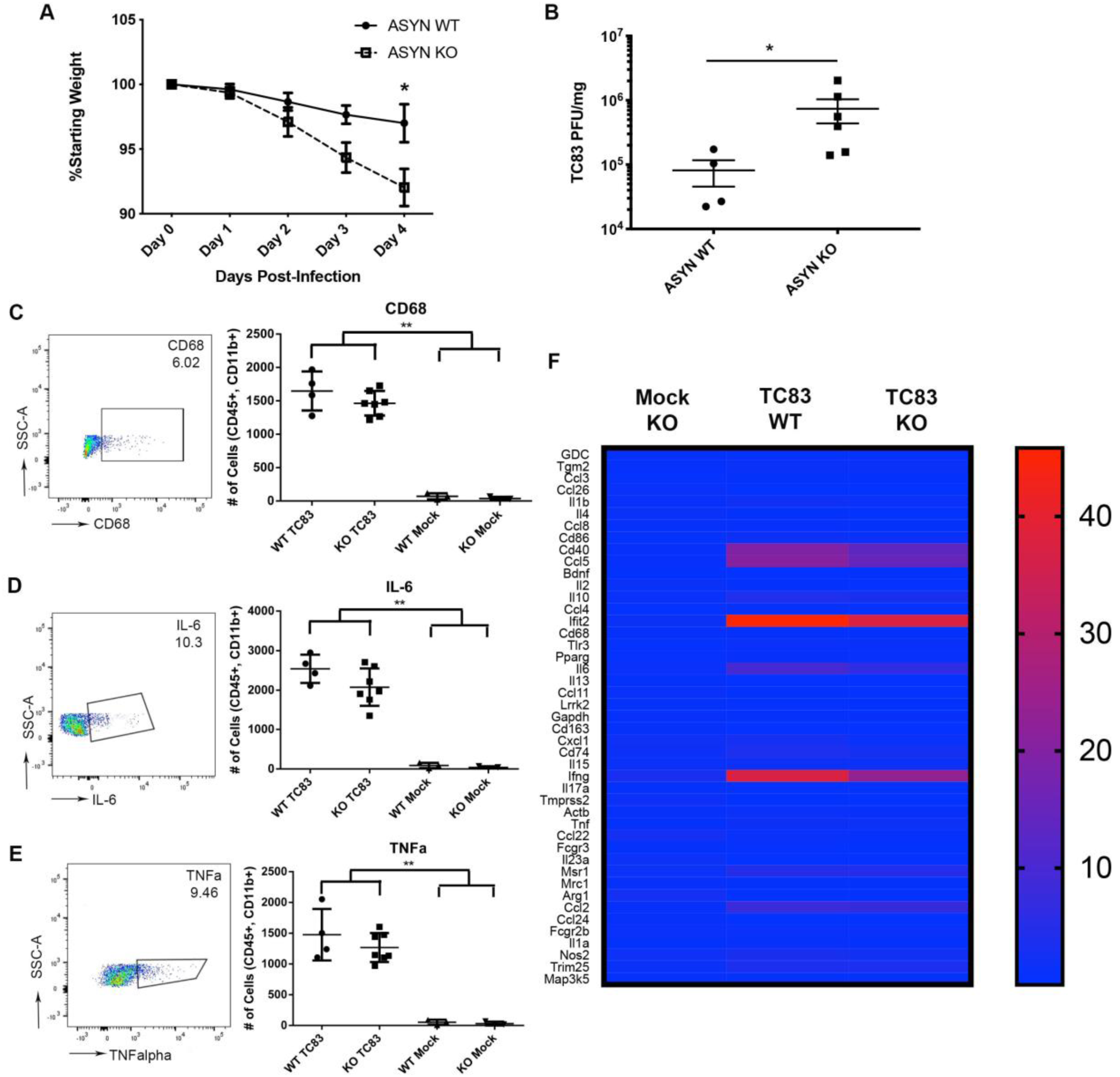
Acute infection with VEEV TC83 results in microglial activation in the brain independent of alpha-synuclein expression. **A**) Asyn knockout (KO) and wild-type (WT) mice were inoculated with VEEV TC83 (10^5^pfu, intracerebral inoculation) and followed for weight loss. *p<0.0001, Two-way ANOVA. N=6 mice per group. **B**) At 4 days post-infection with VEEV TC83, brain tissue from the indicated treatment groups was harvested and analyzed for VEEV TC83 titer using plaque assays. *p=0.038. Mann-Whitney test. N=4-6 mice per group. Asyn KO and WT mice were challenged with TC83 or mock inoculum as above and multiparametric flow cytometry analysis was completed for CD45+CD11b+ cells in the brains of indicated treatment groups for **C**) CD68+, **D**) IL-6, and **E**) TNFalpha expression at day 4 post-infection. Each panel includes a representative flow plot, percent of cells expressing analyte, and a summary graph of flow cytometry data. **p<0.0001. ANOVA, n=4-7 mice per group. **F**) Following mock or VEEV TC83-inoculation by intracranial injection, microglia were isolated from brain tissue and analyzed for interferon stimulated gene expression using PCR array analysis. Data shown with a heat map representation of fold-increase in gene expression from isolated microglia compared to wild-type (WT) mock-inoculated animals. WT=asyn wild-type mice, KO=asyn knockout mice. TC83=VEEV TC83 infection.

### Asyn-dependent expression of ISGs modulates T-cell activation in brain tissue

Previous work has shown that infiltrating CD4+ and CD8+ lymphocytes are critical to control WNV replication in the brain(Shrestha et al., 2006; Sitati and Diamond, 2006; Zhang et al., 2008). Prior studies have also shown that asyn expression supports cytotoxic T-cell responses in Parkinson’s Disease (PD) models(Cebrian et al., 2014; Kannarkat et al., 2013; Seo et al., 2020; Sulzer et al., 2017). When analyzing our RNAseq data, we also found that CD52 and CD274 gene expression was significantly decreased in WNV-infected asyn KO brain tissue compared to virus-infected WT brain tissue. Based on these observations, we evaluated T-cell responses in the brains of virus-infected WT and asyn KO mice using flow cytometry analysis (**Fig. S2**). WT and asyn KO mice were inoculated with mock diluent or WNV (1000pfu, subcutaneous), and brain tissue was collected at day 8 post-infection. Flow cytometry analysis of brain tissue revealed significantly increased CD4+ cells following WNV infection in both WT and asyn KO groups (**Fig. 5**). Since frequencies of T-cells were similar in the brain tissue of WNV-infected WT and asyn KO mice in the brain, we next evaluated activation markers for CD4+ cells. We found that WNV-infected asyn KO mice exhibited significantly decreased frequencies of CD8+CD25+ T-cells, CD4+IFNgamma+ T-cells, TNF+ NK cells, and IL2+ gamma-delta T-cells in the brain tissue compared to WNV-infected WT mice (**Fig. 6**). These data show that asyn-dependent ISG expression in neurons is important to support activation of infiltrating anti-viral T-cell responses in the brain.

**Fig. 5.**
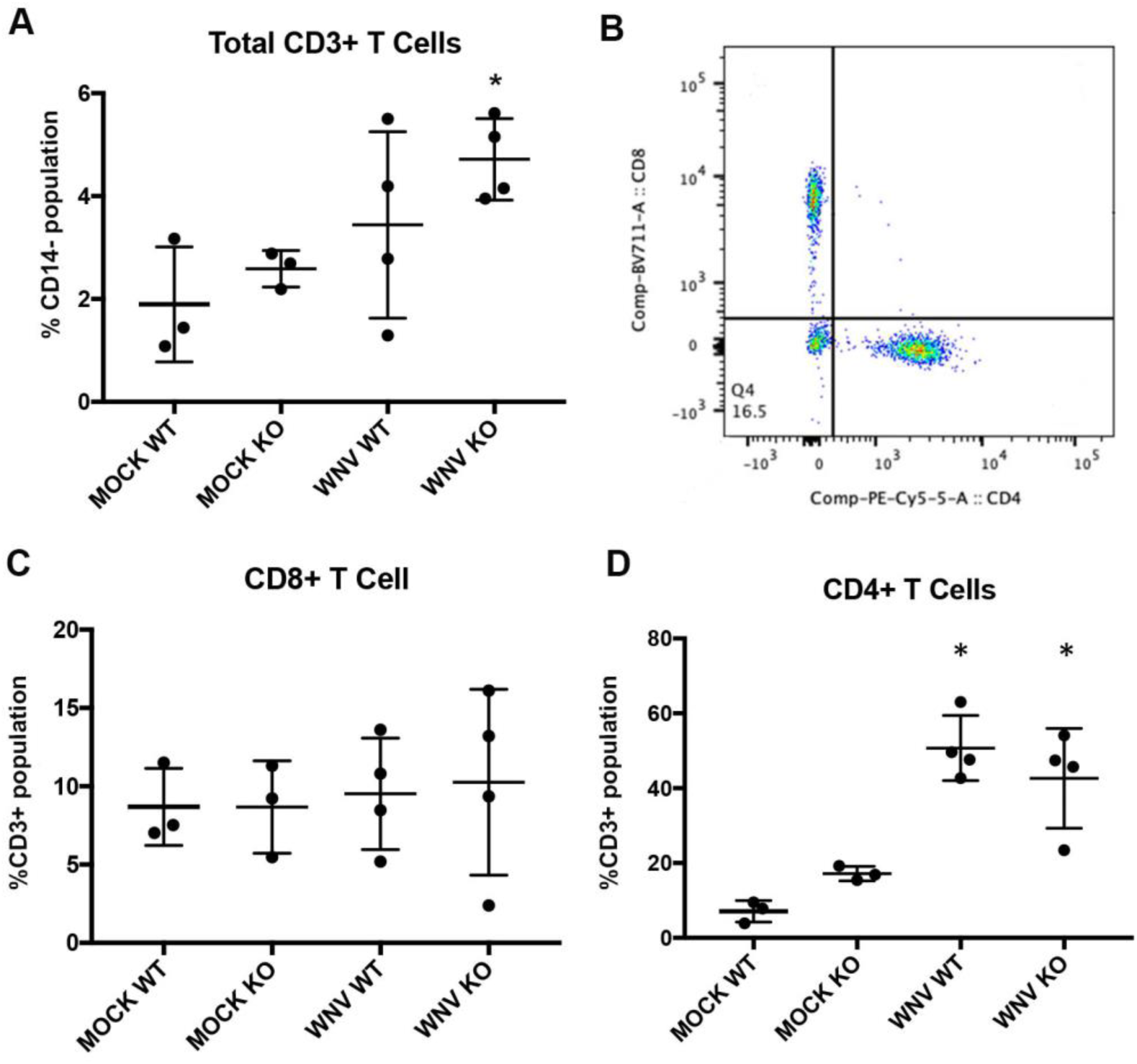
Total T-cell counts in the brain increase to similar degrees in both WT and KO mice following peripheral WNV infection. Mock and WNV inoculated (1000pfu) wild-type (WT) and ASYN KO (KO) mice were sacrificed at day 8 post-infection. Brain tissue was processed for multiparametric flow cytometry for activation markers of T cell subsets based on gating of CD14-cells. **A**) Following WNV infection, CD3+ cells were increased in WNV-infected mice. p<0.05. **B**) Representative gating of CD3+ cells into CD4+ and CD8+ cells following WNV infection in WT mice. **C**) The percentage of CD8+ cells did not significantly change in the brain following WNV infection at day 8 post-infection. **D**) The percentage of CD4+ cells significantly increased following WNV infection in both ASYN WT and KO mouse brain tissue. *p=0.01. ANOVA.

**Fig. 6.**
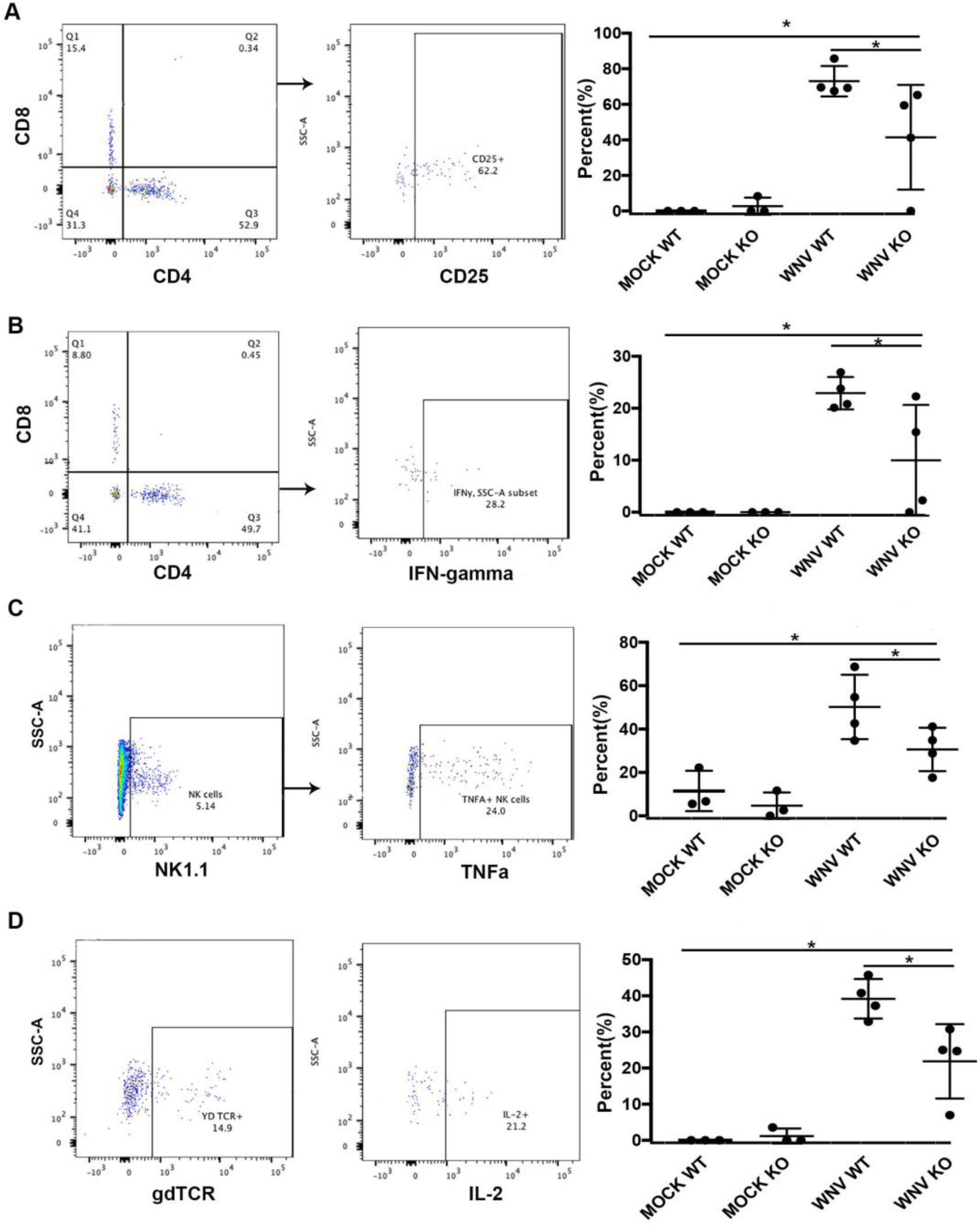
Alpha-synuclein expression supports activation of T-cell effector function in the brain. Asyn WT and KO mice were challenged with mock infection or West Nile virus (WNV, 1000pfu) by subcutaneous footpad inoculation. At day 8 post-infection, animals were sacrificed, T-cells isolated and analyzed by flow cytometry for expression of **A**) CD25+ CD8+ cells, **B**) interferon-gamma+ CD4 T-cells, **C**) TNF-alpha+ NK cells, and **D**) IL2+ gamma-delta (gd)T-cells. *p=0.01. ANOVA, N=3-4 mice per group.

### Acute virus infection increases accumulation of phosphorylated Serine129 asyn in human and non-human primate brain tissue

Our previous data showed that WNV infection increased expression of total asyn in neurons and brain tissue(Beatman et al., 2015). A subsequent study found increased total asyn expression associated with neurons of gastrointestinal tissue following acute viral gastroenteritis(Stolzenberg et al., 2017). Furthermore, a murine model of intranasal inoculated Western Equine Encephalitis virus (WEEV) in mice was able to show increased accumulation of phosphorylated S129 asyn (pS129asyn) in olfactory pathways following infection (Bantle et al., 2019).

Based on these experimental observations, we evaluated histology from human brain tissue obtained at autopsy from patients with acute WNV encephalitis and found evidence of significantly increased levels of pS129+ asyn immunostaining in the grey matter of infected brain tissue compared to non-infected control brain tissue (**Fig. 7A-C**). Due to limited patient numbers, we were unable to comment further on pS129 asyn expression levels for individual anatomical grey matter regions.

**Fig. 7.**
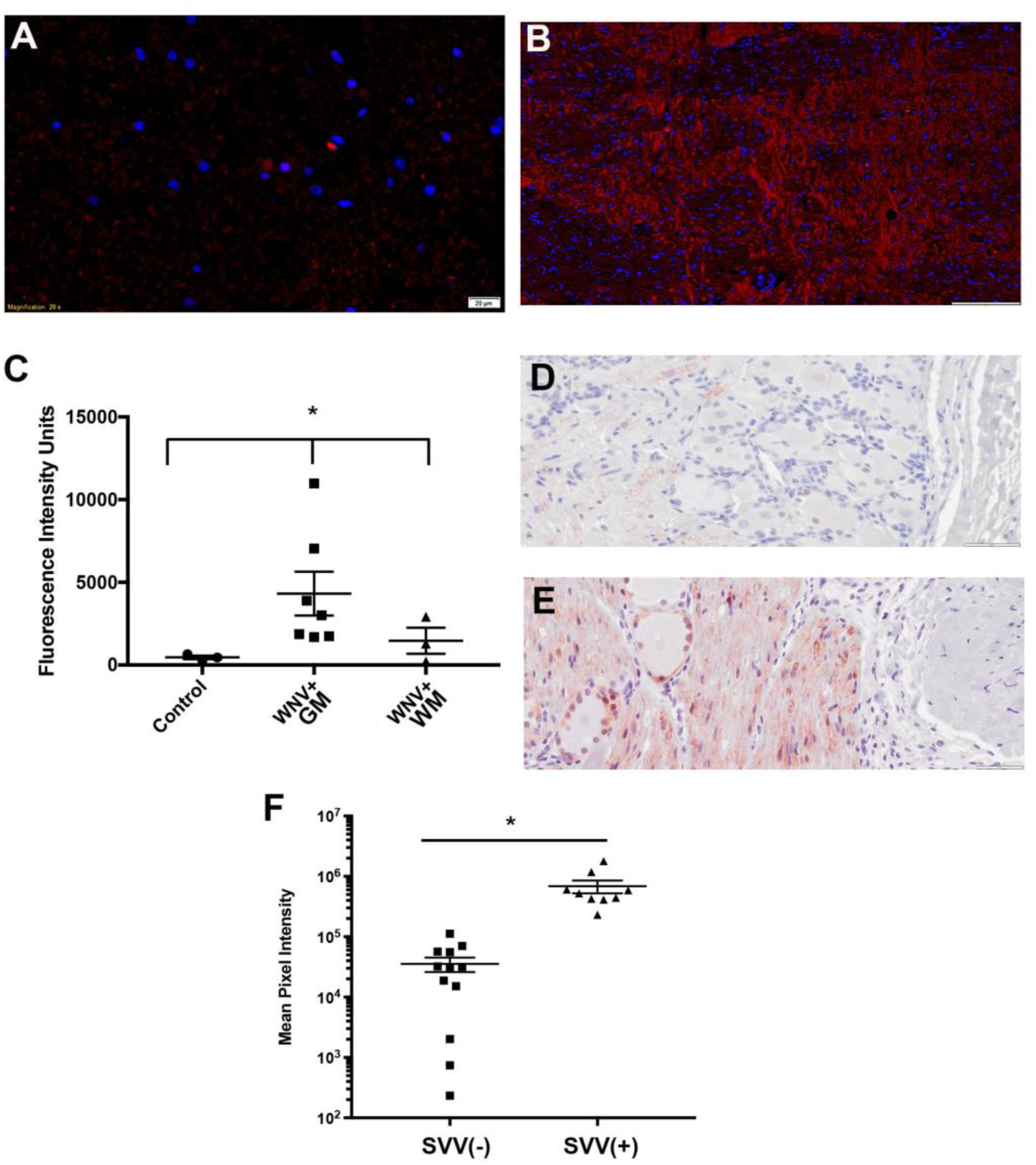
Viral Infection of Human Brain and macaque ganglion results in accumulation of phospho-S129 alpha-synuclein. Immunofluorescent histology labeled tissue for phospho-S129 asyn (red) in **A**) uninfected control human midbrain tissue (bar=20microM). **B**) WNV-infected human pontine nucleus from a patient at day 10 hospitalization with acute encephalitis. Blue=DAPI stain, bar=50mm. **C**) Mean fluorescence intensity units per field of phopho-S129 asyn staining of human brain tissue acutely infected with West Nile virus (WNV). Each data point represents an individual patient in the indicated brain region. *p<0.05 ANOVA. GM=grey matter, WM=white matter. Phospho-S129 asyn IHC (brown) of rhesus macaque lumbar dorsal sensory ganglion from **D)** simian varicella virus (SVV) seronegative control individual and **E**) SVV-reactivated rhesus macaque. Bar=50mm. **F**) Mean pixel intensity of phopho-S129 asyn measured per field from SVV-infected and uninfected rhesus macaque dorsal sensory ganglia. Each data point represents an independent section from 3 individual macaques per group. *p=0.0002, unpaired t-test.

We also evaluated histology of dorsal root ganglia from rhesus macaques after simian varicella virus (SVV), a DNA virus related to human varicella zoster virus, reactivation as previously described(Traina-Dorge et al., 2015). We found increased levels of pS129 asyn in dorsal root ganglion from SVV-infected rhesus macaques compared to seronegative control rhesus macaques (**Fig. 7D-F**). These data indicate that both acute RNA and DNA virus infections induce increased levels of pS129 asyn expression in humans and rhesus macaques, respectively.

### Neuronal spread of pS129asyn pathology is independent of virus-induced microglia activation

Previous studies have shown that peripheral inoculation of WNV causes robust activation of microglia in the brain(Seitz et al., 2018; Vasek et al., 2016). Recent studies have also shown important interactions between neuronal transmission of pathogenic asyn and microglial activation (Grozdanov et al., 2019). Based on these studies, we explored interactions between WNV-induced microglial activation and propagation of pathologic asyn. We first determined if the presence of asyn aggregates in the brain changes the microglial response to WNV infection. We triggered the development of pS129 asyn pathology in WT mice by injection of exogenous asyn pre-formed fibrils (PFFs), or PBS as a control, into the right olfactory bulb. Seven days later, we performed viral challenge using WNV (1000pfu, subcutaneous), and mice were sacrificed at 14 days post-infection for histologic analysis. We found that WNV infection increased microglial activation, as measured by hydraulic radius of Iba1 expressing cells found in the anterior olfactory nuclei (AON) ipsilateral to the injection in the olfactory bulb, to an equal extent in both PBS- and PFF-injected mice (**Fig. 8**). These results indicate that WNV activates microglia independently of the presence of pS129+ asyn pathology in olfactory pathway neurons.

**Fig. 8.**
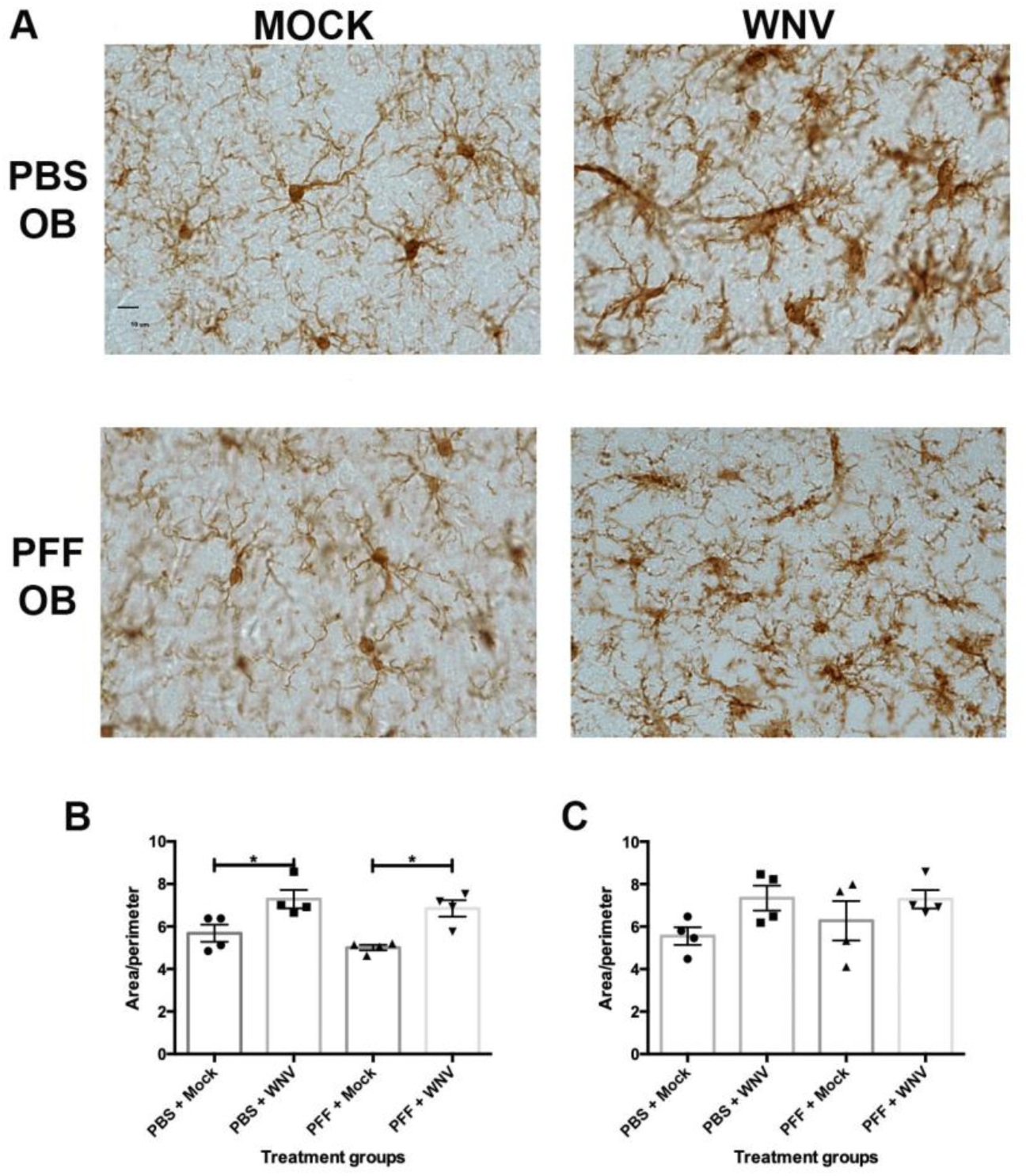
West Nile virus infection increases microglia activation in the brain independent of pre-formed fibril (PFF) treatment. Histological sections of olfactory bulb (OB) were labeled with antisera to Iba1 in mock and WNV inoculated animals. C57B6 mice were inoculated with PFFs or PBS in the right OB and 7 days later, mice were challenged with mock or WNV inoculation (1000PFU, subcutaneous). Brain tissue harvested at day 14 post-infection. **A)** Tissue from each treatment group was labeled with antibody to Iba1 followed by Matlab-guided area:perimeter analysis of microglia in the anterior olfactory nuclei, **B)** ipsilateral and **C)** contralateral to PFF injection. *p<0.05, ANOVA. AON=anterior olfactory nucleus.

Since the WNV-infection leads to upregulation of asyn expression, we also explored the role of WNV-induced upregulation of asyn expression and microglial activation on the development of pS129 asyn pathology in olfactory pathways (**Fig. S3A**). The same treatment groups as above were sacrificed at an acute (2 weeks) and chronic (11 weeks) time point following WNV challenge and brain tissue was analyzed and scored for pS129+ asyn pathology (**Fig. S3B**). Following histologic scoring of both the ipsilateral and contralateral AON, we found no significant difference in pS129+ asyn pathology scores at any of the time points following analysis of AON tissue at the ipsilateral and contralateral regions from OB injection of PFFs (**Fig. 9**). These data suggest that a single exposure to neuroinvasive virus does not significantly alter levels of pS129+ asyn pathology triggered in this murine model.

**Fig. 9.**
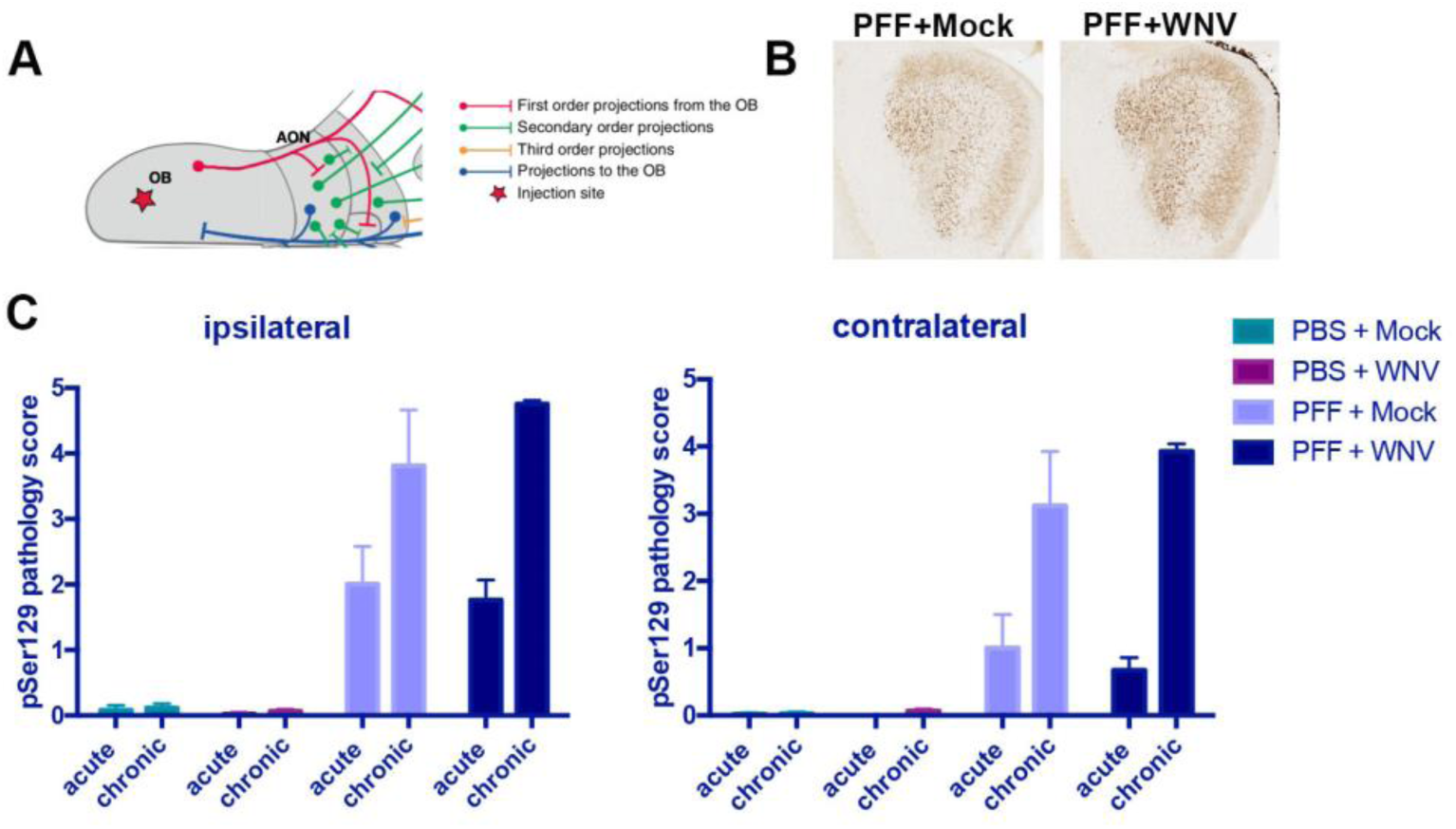
Single exposure to peripheral West Nile virus infection does not significantly increase PFF expression in the Anterior Olfactory Nucleus (AON). WT mice were challenged with mock or WNV (1000pfu) one week following OB inoculation of PBS or pre-formed fibrils (PFFs) and sacrificed at 2 weeks (acute) and 11 weeks (chronic) post-infection to (**A**) evaluate expression of phospho-S129 asyn in the anterior olfactory nucleus (AON). **B**) Representative histology image of anti-phospho-S129 asyn (brown) in the AON. **C**) Summary data of phospho-S129 asyn scoring in the AON in the indicated treatment groups. N=3-4 mice per group.

## Discussion

Asyn is a highly conserved protein expressed in neurons of most vertebrate species. Its unique localization to nuclei as well as synaptic puncta suggest it may have functions beyond synaptic biology. Mounting evidence supports an important role for asyn in protection of neurons but the mechanisms of protection remain unclear (Beatman et al., 2015; Schaser et al., 2019). Our data now show that during virus infection, asyn supports expression of ISGs required to inhibit viral infection. Moreover, the same ISGs stimulated by virus infection are also dependent on asyn for expression following poly I:C and type I interferon treatment. The importance of asyn-dependent ISG expression is supported by our findings showing that TC83 growth is significantly increased in asyn KO neurons and in asyn KO brain tissue compared to WT neurons and mice, respectively. TC83 is attenuated due to a mutation in a 5’ untranslated region that results in increased IFIT1 restriction of viral growth(Reynaud et al., 2015), and our discovery of increased viral growth in the absence of asyn suggested that IFIT1 expression was decreased in asyn KO neurons. We subsequently found evidence of decreased IFIT1 expression in asyn KO neurons following TC83 infection and following treatment with Poly I:C. These data show that asyn expression supports of ISG responses that are critical to control RNA virus infection in neurons.

We also found that TC83 infection and poly I:C treatment did not significantly increase expression of *TRIM25* in WT human neurons, while treatment with type 1 interferon resulted in significantly increased expression of *TRIM25*. In human asyn KO neurons, expression of TRIM25 was significantly decreased compared to WT neurons following treatment with type 1 interferon. Since type 1 interferon is down-stream of poly I:C mediated activation of innate immune responses and asyn-dependent expression of *TRIM25* occurs down stream of type 1 interferon stimulation but not poly I:C stimulation, we hypothesize that asyn interacts with type 1 interferon signaling pathways to support and regulate ISG expression in neurons.

We next evaluated the role of asyn-dependent neuronal expression on activation of other immune cell responses in the brain. Since asyn expression is known to activate microglia, we evaluated the role of asyn-dependent ISG expression on microglia activation following both WNV and TC83 infection. We found that microglia activation was significantly increased following virus infection but was unaltered in the presence or absence of asyn expression in the brain. These data show that the innate immune role of microglia in the brain during viral infection is independent of asyn expression.

Since T-cell responses are critical to clear WNV from brain tissue, we determined the role of asyn expression on infiltrating lymphocytes in the brain(Brien et al., 2009; Brien et al., 2008; Glass et al., 2005; McCandless et al., 2008; Shrestha and Diamond, 2004; Sitati and Diamond, 2006). We found that infiltrating CD4+ T-cells exhibited significantly decreased activation markers in asyn KO brain tissue following WNV infection. We also evaluated NK cell and gamma-delta T-cell responses as anti-viral lymphocytes with activity independent of MHC interactions. We again found significantly decreased activation of infiltrating lymphocytes in brain tissue of asyn KO mice following WNV infection. These data show that asyn expression in the brain supports activation of infiltrating lymphocytes.

Asyn is believed to play a role in the pathogenesis of PD and evaluating links between innate immune signaling in PD are important to fully define the pathogenesis of PD. Accumulating evidence indicates that specific triggers can upregulate expression and cause post-translational modifications of asyn such as phosphorylation, glycation, or truncation(Bantle et al., 2019; Stolzenberg et al., 2017; Vicente Miranda et al., 2017). These changes are believed to promote the formation of Lewy pathology, which is thought to be key to the development and progression of PD. Our studies show that virus infection in humans and non-human primates resulted in increased levels of phosphorylated S129 asyn in neurons. These data show that acute virus infection triggers asyn modifications found in PD.

We next determined if a single exposure to virus enhances spreading pathology of pS129 asyn in the murine model that was previously described(Rey et al., 2016). Following olfactory bulb injection of preformed fibrils, we challenged mice with peripheral inoculation of WNV. While WNV infection resulted in significantly increased microglia activation, we found no evidence that pS129 asyn pathology was enhanced following a single virus challenge. These data show that acute activation of microglia by inflammatory responses or the upregulation of asyn expression are not sufficient to increase levels of pS129 asyn accumulation in this experimental model. Further studies are needed to fully evaluate the interactions between innate immune activation and chronic spread of pS129 asyn pathology.

Based on our data, we propose that specific innate immune triggers increase expression of asyn and failure to degrade up-regulated or modified asyn may increase the risk for PD. If fundamental innate immune interactions drive asyn misfolding and subsequent development of Lewy pathology, then inhibition of these interactions may provide a novel target for therapies that reduce the risk for PD developing in the first place. Further studies evaluating the interactions between innate immune signaling and asyn proteostasis should reveal new insights into the pathogenesis of disease processes associated with asyn expression.

## Materials and Methods

### Ethics statement

All animal work was performed at the University of Colorado Anschutz Medical Campus in accordance to and following approval by the Institutional Animal Care and Use Committee. All work with live viruses and recombinant DNA was approved by the University of Colorado Institutional Biosafety Committee and performed in accordance with local and national regulations of pathogens. Human brain tissue was obtained from de-identified human autopsies at the University of Colorado Hospital with approval for non-human research by the local Colorado Multiple Institutional Review Board. All work with hESCs was completed at the University of Edinburgh and ethics approval was granted by the MRC Steering Committee for the UK Stem Cell Bank and for the Use of Stem Cell Lines (ref. SCSC13-19). Differentiated neurons were provided to other laboratories for use following gene deletion and maturation. Support for hESC work was supported by grants through the Kunath laboratory.

### Cell culture

All cell lines were maintained at 37°C in 5% CO2. Vero (ATCC CCL81) and BHK-21 cells were maintained in minimum essential medium containing Earle’s salts and L-glutamine supplemented with 1% penicillin-streptomycin, 10% heat-inactivated fetal bovine serum, 1% nonessential amino acids, and 1% sodium pyruvate.

*Snca*^*-/-*^ and *Snca*^+/+^ human embryonic stem cells were generated and differentiated towards midbrain dopaminergic neuron for 16 by the Kunath laboratory as previously described(Chen et al., 2019). Following this, the cells were frozen and shipped to the Beckham laboratory. Cells were then thawed and plated in L111-coated 48-well plates at a density of 800,000 cells/cm2. Cells were differentiated for the following 26 days (42 days of total differentiation) in neuronal differentiation media consisting of Neurobasal Media (Thermo Fisher Scientific) + B27 supplement (without Vitamin A, 1:50, Thermo Fisher Scientific) + l-Glutamine (2 mM, Thermo Fisher Scientific) supplemented with ascorbic acid (AA, 0.2 mM, Sigma), brain-derived neurotrophic factor (BDNF, 20 ng/ml, Peprotech), glial cell line-derived neurotrophic factor (GDNF, 10 ng/ml, Peprotech), dibutyryl cyclic AMP (dcAMP, 0.5 mM, Sigma), and DAPT (1 μM, Tocris). Y27632 (Y2, 10 μM, Tocris) was present in medium from day 16 to day 17. Cellular media was removed and replaced every 3-4 days.

### Virus propagation and quantification

West Nile virus strain 385-99 (NY99) was obtained from clone derived virus and propagated in *Aedes albopictus* (C6/36, ATCC CRL-1660) cells as previously described(Beatman et al., 2015). Venezuelan Equine Encephalitis virus (VEEV) TC83 isolates were obtained from the laboratory of Dr. Michael Diamond at Washington University in St. Louis and was propagated in BHK cells. Herpes Simplex virus type I (HSV-1) strain F was procured from ATCC (#VR733) and passaged over *Cercopithecus aethiops* kidney cells (Vero cells, ATCC CCL-81) at 34 °C. Viral titers for all viruses were quantified in Vero cells by standard plaque assay as previously described(Beatman et al., 2012). Viral genome was quantified by probe-based qRT PCR.

The 3’UTR of WNV was amplified using forward primer CAG ACC ACG CTA CGG CG, reverse primer CTA GGG CCG CGT GGG, and probe /6FAM/TCT GCG GAG AGT GCA GTC TGC GAT/MGBNFQ/. The UL30 gene of HSV-1 was primed using forward primer CGC GTC CAA GCC CCG CAA, and reverse primer GGT GCC ACA CTT CGG GAA TAA ACC T, and quantified using probe /6FAM/ CCT CGG CCA GCT CGG ACA CCA /GMGBNFQ/. The nsP1 gene of VEEV TC83 was amplified using the forward primer GCC TGT ATG GGA AGC CTT CA, reverse primer TCT GTC ACT TTG CAG CAC AAG AAT, and probe 6-FAM/ CCT CGC GGT /ZEN/ GCA TCG TAG CAG C/ 3IABkFQ/. Quantification was achieved by generating standard curves of serial diluted plasmids of known copy number and normalized to 18s rRNA copies. For 18s rRNA quantification, priming was achieved with the forward primer CGC CGC TAG AGG TGA AAT TC, reverse primer sequence CAT TCT TGG CAA ATG CTT TCG, and probe /6-FAM/CAA GAC GGA CCA GAG CGA AAG CAT/TAMRA/.

### Mouse and Primate Studies

*Snca*^*-/-*^ mice were obtained from Jackson Laboratories (#3692) and back-crossed seven generations to C57B/6J mice (#664). Microsatellite analysis performed by Jackson Laboratories confirmed mice were 96.3% C57B/6J. These mice were crossed with WT C57B/6J mice to generate *Snca*^*+/-*^ heterozygous mice. Genotyping by conventional PCR was routinely performed to confirm *Snca* status as described.(Beatman et al., 2015) Mice transgenic for human alpha-synuclein and deficient for murine alpha-synuclein (Tg*SNCA*^WT^;*Snca*^*-/-*^) were obtained from Jackson Laboratories (#10710) and crossed with mice heterozygous for murine alpha synuclein (*Snca*^*+/-*^). The offspring of these mice were tested via conventional PCR for the presence of both the human and murine alpha-synuclein genes to generate mice that had only the transgenic human alpha-synuclein (Tg*SNCA*^WT^;*Snca*^*-/-*^) and mice that had both the transgene and were heterozygous for murine alpha-synuclein (Tg*SNCA*^WT^;*Snca*^*+/-*^). Virus used for infections was first diluted to the appropriate viral titer in HBSS before being administered by subcutaneous injection, intracranial injection, or corneal inoculation.

Prior to subcutaneous inoculation of virus, all mice were randomized, weighed, and placed under isoflurane-induced anesthesia. Equal numbers of male and female mice were used for all studies with the exception of the use of only female mice for the total brain RNAseq analysis. For subcutaneous injections, 10 uL of virus solution was injected into the left footpad of mice with the use of a Hamilton syringe. For intracranial injections, 10 uL of virus solution was injected 3 mm deep into the brain near the Bregma with the use of a 25 gauge needle. Mice were monitored for morbidity and weighed daily. For HSV-1 inoculations, the corneas of anesthetized mice were scarred using a 31 gauge needle 9 times using a crosshatch pattern. Subsequently, 10^5^ PFU HSV-1 was administered to the eye in 10 uL via pipette. Mice losing more than 15% bodyweight prior to the end of the study were euthanized and excluded from the study for humane reasons.

At the end of each experiment, mice were euthanized by isoflurane overdose before proceeding with tissue harvest. All mice were perfused with 20 mL of phosphate buffered saline solution (PBS) prior to tissue harvest. In the case of histology analysis, intracardiac perfusion of 4%PFA was completed. Samples collected for RNA gene expression assays, immunostaining assays, or viral quantification were collected and stored in RNALater (Invitrogen, #AM7021), 10% neutral buffered formalin (NBF), or PBS, respectively. Samples stored in RNALater and PBS were stored at -80 C until needed. Samples fixed in 10% NBF were stored at room temperature for 48 hours and subsequent storage at 4 °C in PBS until processed for paraffin embedding. Plasma was collected at time of euthanasia by cardiac stick, collected in EDTA coated microtubes. Blood cells were pelleted by centrifugation at 1500 rcf for 5 minutes at 4 °C, aliquoted and stored at -80 C until needed.

#### Unilateral injections of asyn fibrils into the olfactory bulb

Mouse α-synuclein amyloid aggregates were produced and kindly provided by Dr. Jiyan Ma, Van Andel Institute, USA. Before surgery, asyn fibrils were produced by the sonication of α-synuclein amyloid aggregates in a water-bath sonicator for 10 min and were maintained at room temperature until injection. Mice were anesthetized with isoflurane and injected unilaterally in the right OB with either 0.8 µL of asyn fibrils (5 µg/µl) or 0.8 µL of PBS (phosphate buffered saline), as a control (coordinates from bregma: AP: + 5.4 mm; ML: - 0.75 mm and DV: -1.0 mm from dura). Injections were made at a rate of 0.2 µL/min using a glass capillary attached to a 10 µL Hamilton syringe. After injection, the capillary was left in place for three min before being slowly removed. Prior to incision, the animals were injected with local anesthetic into the site of the future wound margin (0.03 mL of 0.5% Ropivacaine;

Henry Schein, USA). Following surgery mice received 0.5 mL saline s.c. for hydration.

#### Primate Studies

Simian varicella virus infection of non-human primates and collection of tissue samples were performed in the Tulane National Primate Research Center (TNPRC) in accordance with the recommendations of the US Department of Agriculture Animal Welfare Act regulations, the Guide for the Care and Use of Laboratory Animals, and the Institutional Animal Care and Use Committee (IACUC) at Tulane University and the TNPRC. The protocol for SVV infection and tissue analysis was approved ty the IACUC of Tulane University and the TNPRC.

### Brain digestion and cell isolation

Cells collected from adult, infected mouse brain for flow cytometry were obtained as described.(Posel et al., 2016) Isolated cells were incubated overnight (5 hours for microglia analysis) in GolgiPlug and GolgiStop (BD Bioscience #555029 and #554724) in RPMI media containing HEPES and 10% fetal bovine serum (FBS) prior to staining and analysis by flow cytometry. Microglia were isolated from adult mouse brain using the multi tissue dissociation kit 1 (Miltenyi Biotec #130-110-201), followed by CD11b microglia isolation kit (Miltenyi Biotec #130-093-634) according to manufacturer’s protocols. Isolated cells were immediately stored in TRK lysis buffer (Omega) containing 25 uL/mL beta mercaptoethanol. Cells were stored in lysis buffer at -80°C prior to RNA extraction.

### Flow cytometry

The following antibodies were used for extracellular flow cytometry: anti-mouse CD45 BV650 (clone 30-F11, Biolegend), anti-mouse/human CD11b APC-Cy7 (clone M1/70, Biolegend), anti-mouse CD11c PE-eFluor 610 (clone N418, eBioscience), anti-mouse CD103 BV711 (clone 2E7, Biolegend), anti-mouse Ly6C BV785 (clone HK1.4, Biolegend), anti-mouse CD14 BV605, anti-mouse CD19 BV605, anti-mouse γδ TCR BV421, anti-mouse CD25 BV650, anti-mouse CD8 BV711, anti-mouse CD4 PE-Cy5.5, anti-mouse NK1.1 PE-CF 594, anti-mouse CD68 PE-Cy7, and anti-mouse CD3 BV785 (clone 17A2, Biolegend). The following antibodies were used for intracellular flow cytometry: anti-mouse CD68 PE-Cy7 (clone FA-11, Biolegend), anti-mouse IL-17 APC, anti-mouse IL-2 APC-Cy7, anti-mouse TNFα PE-Cy7, anti-mouse TNFα BV421,, anti-mouse IFNγ PE, anti-mouse IFNγ AF700, anti-mouse TNFa AF700 (clone MP6-XT22, Biolegend), anti-mouse IL-6 APC (clone MP5-20F3, BD Pharmingen), and anti-mouse IL-12 (p40/p70) PE (clone C15.6, BD Biosciences). Ghost Violet 510 dye (Tonbo Biosciences) was used to assess viability.

For flow staining, single cell suspensions were washed in PBS, centrifuged at 500 rcf for 5 minutes, and briefly vortexed. Next, 10 µL of viability dye (0.1 µL dye + 10 uL FACS buffer per sample) was added to each sample, vortexed, and incubated at room temperature for 10 minutes. Next, 50 µL of extracellular antibodies prepared in FACS buffer were added directly to each sample, vortexed, and incubated at 4°C for 25 minutes. 210 µL of Cytofix/Cytoperm solution (BD Biosciences) per sample was then added to permeabilize the cells, followed by vortexing and incubation for 20 minutes at 4°C. The cells were then washed in 1 mL of 1x Perm/Wash buffer (BD) or Flow Cytometry Perm Buffer (Tonbo Biosciences) twice, centrifuged at 700 rcf for 5 minutes, and vortexed. Next, 50 µL of intracellular antibodies in Perm/Wash or Perm buffer were added to each sample, vortexed, and incubated at 4°C of 45 minutes. The samples were then washed once more in Perm/Wash buffer, centrifuged at 700 rcf for 5 minutes, vortexed, and finally fixed in 1% paraformaldehyde (Thermo Fisher).

The data was acquired on a LSRII flow cytometer (BD) using voltages standardized according to previously published methods (Perfetto et al., 2012). FlowJo software (FlowJo, LLC, Ashland, Oregon) was used to analyze the data. The gating strategies used are shown for individual experiments in the supplemental figures.

### Cell Culture

*Snca*^*-/-*^ and *Snca*^*+/+*^ hESC derived midbrain dopaminergic neurons were grown as described above. Following this, the cellular media was removed and replaced with complete neuronal differentiation media (described above) containing VEEV-TC83 virus at a multiplicity of infection (MOI) of 1. 300 ul of the media was removed and replaced with fresh, virus-free neuronal differentiation media every 12 hours for 72 hours. The viral content of these samples was then titered via plaque assay as described previously(Beatman et al., 2015). Differentiated neurons were infected with VEEV-TC83 at an MOI of 10. 12 hours post-infection, RNA from these cells was extracted and analyzed via qPCR (described below). Differentiated human primary neurons were treated with 10,000 IU/mL of mixed-type human IFNa, 25µg/mL of Poly(I:C), or mock treated. RNA from these cells was extracted 4 hours post-treatment and analyzed via qPCR.

### Gene expression assays

RNAs collected from tissue were extracted using Trizol-chloroform extraction followed by column based isolation using the E.Z.N.A Total RNA kit II (Omega Bio-Tek #R6934) according to manufacturer’s protocol. RNAs collected from cells were extracted using E.Z.N.A Total RNA kit I (Omega Bio-Tek #R6834) according to manufacturer’s protocol. Qiagen’s RT_2_ custom PCR array was used to assay isolated microglia and neurons from infected mouse brain for interferon stimulated genes and microglia activation markers. RNA extraction and purification, cDNA synthesis, and Qiagen RT_2_ custom PCR array was performed according to manufacturer’s protocol and recommendations.

BioRad’s PrimePCR probe-based qPCR assays for Oas1b, IRF9, TLR3, Trim25, TGTP1, and MAP3K were used to verify RNAseq results. RNA extraction, cDNA synthesis, and qPCR was performed according to manufacturer’s protocols and recommendations. Normalization was achieved using 18s rRNA copy number as described above. BioRad’s PrimePCR SYBR Green-based qPCR assays for *OAS1b, TLR3, TRIM25*, and *IFIT1* were used to analyze gene expression during infection and following immune-pathway activator treatment (e.g. Poly I:C) of differentiated *SNCA*^-/-^ and *SNCA*^+/+^ hESC derived neurons. RNA extraction, cDNA synthesis, and qPCR were performed according to manufacturer’s protocols and recommendations. Normalization was achieved by calculating the ΔCT value of each sample compared to expression of the housekeeping gene GAPDH. Relative expression was calculated by comparing the ΔCT value of each sample to the average ΔCT value of the mock infected, WT mouse samples.

RNAseq analysis was performed on bulk brain RNAs by Novogene (Sacramento, CA, USA). Analysis of sequencing reads completed by Novogene and additional analysis of Fastq files completed in the Beckham laboratory. Following total RNA quantification (Nanodrop) from brain tissue, we completed mRNA enrichment(poly-T oligo-attached magnetic beads), cDNA synthesis, end repair, poly-A and adaptor addition, fragment selection and PCR, and library quality assessment (Agilent2100) followed by Illumina sequencing (NovaSeq). Following data clean up, clean reads representing 96.51% of total reads were available for analysis at approximately 60-80million reads per sample. Mouse genome sequence alignment was completed with STAR with 85.7% of reads mapping to exons and fragments per kilobase of transcript sequence per millions base pairs sequenced (FPKM) calculated for gene expression. Overall gene expression was analyzed using principal component analysis and DESeq2 R package for differential expression analysis. ClusterProfiler software was used for enrichment analysis including GO enrichment, DO enrichment, KEGG and Reactome database enrichment.

### Immunostaining assays

Five mm thick sections from formalin-fixed, paraffin-embedded tissue was used for immunostaining assays. The following primary antibodies were used; rabbit monoclonal antibody to serine 129 phosphorylated α-synuclein (1:200, Cell Signaling Technologies #23706), guinea pig antiserum to Iba-1 (1:100, SySY #234-004), chicken polyclonal antibody to GFAP (1:50, Abcam #ab4674), rabbit polyclonal antibody to WNV envelope protein (1:800, Novus Biologicals #NB100-56744), and rabbit polyclonal antibody to Iba-1 (1:200, Wako 019-19741). For immunohistochemistry, rabbit primary antibodies were detected with ImmPRESS HRP anti-rabbit IgG polymer detection kit (MP-7401) and ImmPACT NovaRED peroxidase substrate kit (SK-4805) according to manufacturer’s protocol. For immunofluorescence the following detection antibodies were used; Cy5-conjugated goat anti rabbit IgG (1:200, Jackson ImmunoResearch #111-175-144), TRITC-conjugated goat anti chicken IgY (1:200, Jackson ImmunoResearch #103-025-155), and FITC-conjugated goat anti guinea pig IgG (1:200, Rockland #106-102-002). Images were acquired using an Olympus VS120 slide scanning microscope and analyzed using ImageJ.

For the WNV study combined with asyn fibrils, 40 micron free-floating brain sections were incubated with 3% H_2_O_2_ for 20 min to quench endogenous peroxidase activity, blocked for 1 h with 5% normal goat serum and 0.3% Triton X-100, then incubated overnight at room temperature with Iba1 primary antibody (1:800, WAKO). Sections were incubated with goat anti-rabbit biotinylated secondary antibody (1:500, Vector Laboratories) followed by incubation with Avidin/Biotin ABC reagent (Vector Laboratories). Immunolabeling was revealed using diaminobenzidine, which yielded a brown-colored stain visible in bright-field light microscopy. Images were captured with a color Retiga Exi digital CCD camera (QImaging) using NIS Elements AR 4.00.08 software (Nikon) using the 60x magnification (oil immersion, 1.40 N.A.) on a Nikon Eclipse Ni-U microscope (Nikon). We assessed the morphology of microglia to determine to state of activation as activated microglia typically have a larger area:perimeter ratio. To do this we processed RGB color images using a custom MATLAB (Mathworks) script.

### Immunoglobulin ELISAs

Total IgG and IgM from *Snca*^*+/+*^ and *Snca*^*-/-*^ were measured by commercially available ELISA kits (Invitrogen, #88-50400-22 and #88-50470-22) according to manufacturer’s protocols. For WNV specific IgG, WNV particles were purified by ultracentrifugation over a 20% sucrose buffer, washed once and stored in PBS. Standard BCA assay was used to measure protein concentration (Pierce #23227). Two hundred and fifty ng whole WNV particles were plated in 100 uL of PBS into 96-well ELISA compatible microplates overnight at 4 °C. Plates were blocked in 5% BSA for 2 hours at room temperature before applying mouse plasma diluted in PBS for one hour at 1 hour at room temperature. The mouse monoclonal pan-flavivirus antibody 4-G2 was used as a positive control and for generating standard curves. HRP-conjugated Donkey anti mouse IgG (1:600, Jackson ImmunoResearch #715-035-150) followed by TMB substrate were used for detection.

### Statistical analysis

All statistical studies were completed using Gaphpad Prism software with the indicated statistical test in the text.

## Acknowledgments

We thank Dr. Vicki Traina-Dorge who performed the SVV inoculation and collection of tissue samples for prior work and allowed us to utilize samples from prior studies. We thank Emily Schulz for her technical assistance in the asyn fibril injections. This work was supported by VA Merit Funding (I01BX003863) and DOD PRMRP funding (contract W81XWH-17-1-0183) to J.D.B. M.E.J., J.A.S., M.L.E.G and P.B. are supported by a grant from Farmer Family Foundation. Y.C. and T.K. were supported by funding from the UK Centre for Mammalian Synthetic Biology.

## Author contributions

A.M. completed RNAseq studies, brain gene expression studies, microglia isolations and gene expression, and histology/imaging analysis. B.M. completed all experiments with primary neurons, microglial flow cytometry, and completed all experiments with VEEV TC83. Y.C. and T.K. created *SNCA* gene deletion hESCs and human neurons and provided support for human neuronal experiments. M.E.J., J.A.S., M.L.E.G. and P.B. provided training, support, and analysis of brain tissue in experiments with injected fibrils. J.M. created and supplied PFFs. B.K.D. provided human brain tissue samples and guided human autopsy studies. R.M. provided primate ganglion samples for analysis from macaque SVV studies. J.D.B. designed the studies, guided and completed all data analysis, synthesized all data, and wrote the manuscript with input from all co-authors.

## Data and materials availability

All materials are available upon reasonable request following materials transfer agreement. All data is available in the main text or the supplementary materials, and all sequence data is available through NCBI.

## Figures and Legends

**Fig. S1.**
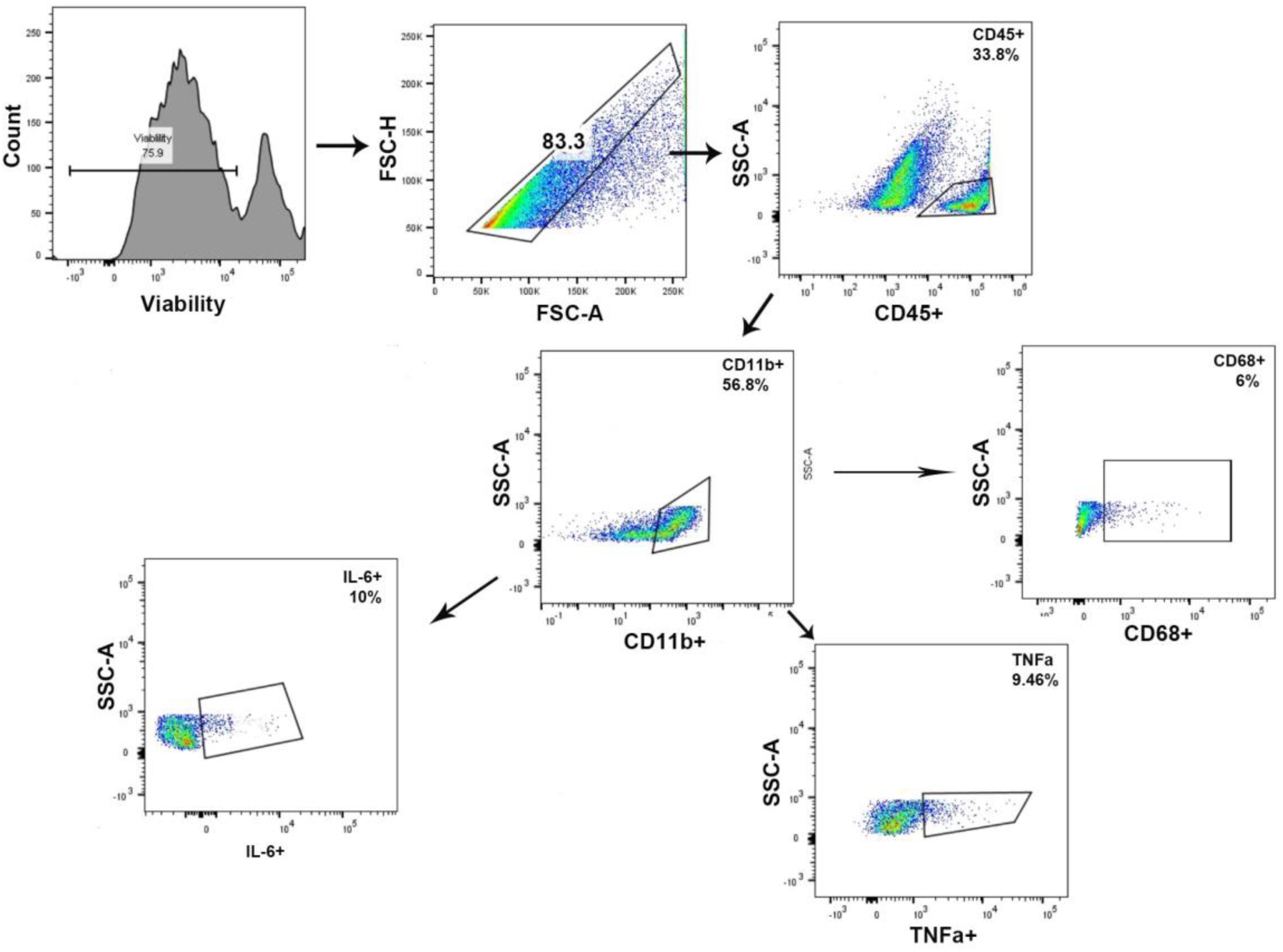
Gating strategy for flow cytometry analysis of CD45+ cells in the brain. Cells were gated on viability and singlets were analyzed for CD45+ expression. Of the CD45+ cells, CD11b+ cells were analyzed for expression of CD68, TNFalpha, and IL-6.

**Fig. S2.**
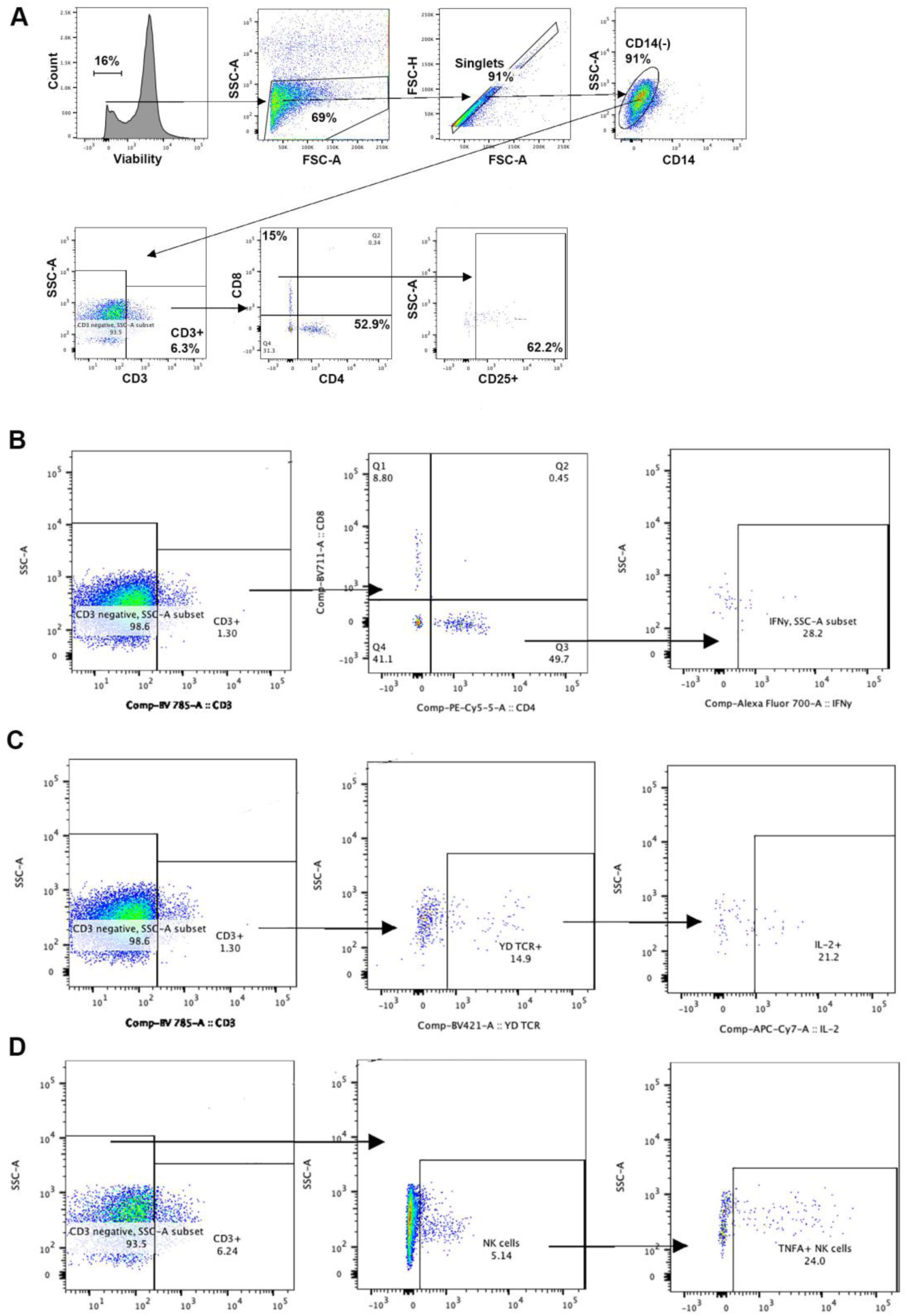
Gating strategy for flow cytometry analysis of T-cells in the brain. Following viability selection, single cells were evaluated for CD14 expression. CD14-cells were analyzed for CD3 expression and CD3+ cells were gated for CD4+ or CD8+ expression followed by determination of activation markers including **A**) CD25, **B**) CD4+, IFN-gamma, **C**) CD3+ gdTCR+ IL2+, **D**) CD3+ NK1.1+ TNFalpha+.

**Fig. S3.**
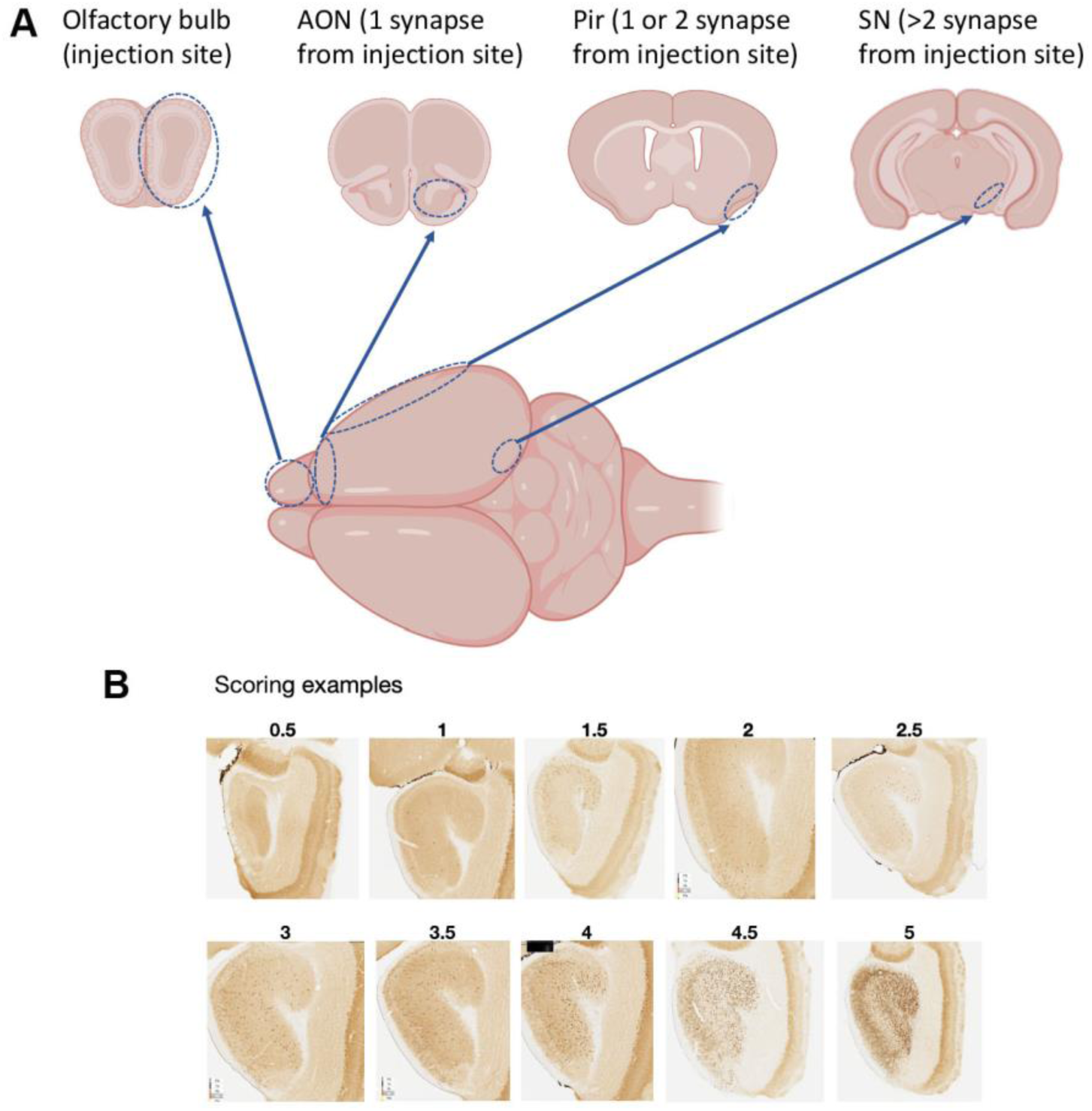
Phospho-S129 asyn Scoring System in Murine brain tissue. **A**) Diagram showing anatomic localization of brain regions evaluated during PFF inoculation and spread. **B**) Histologic images showing phospho-S129 asyn-positive sections indicative of specific scores from 0.5 to 5.

**Table 1.**
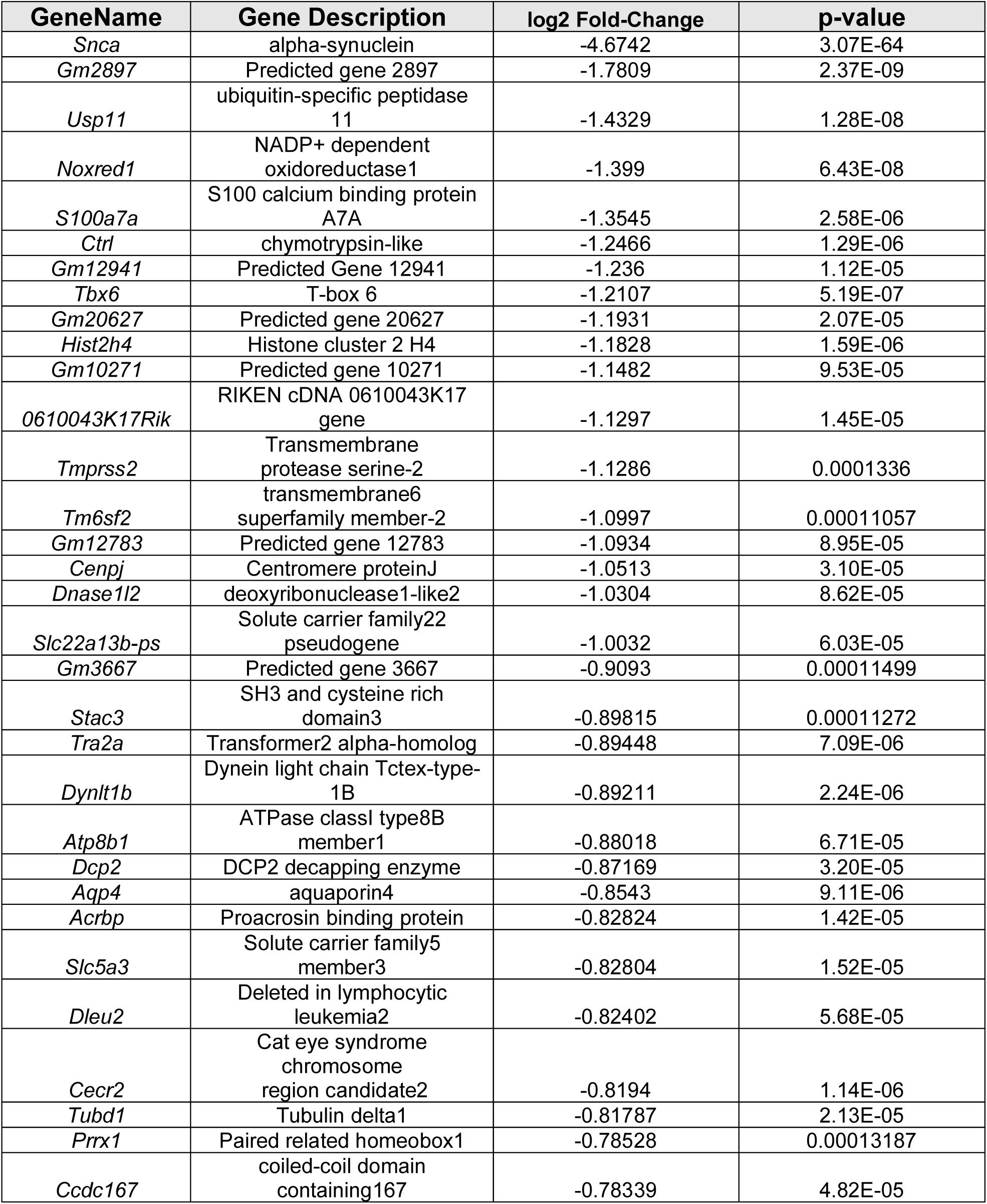
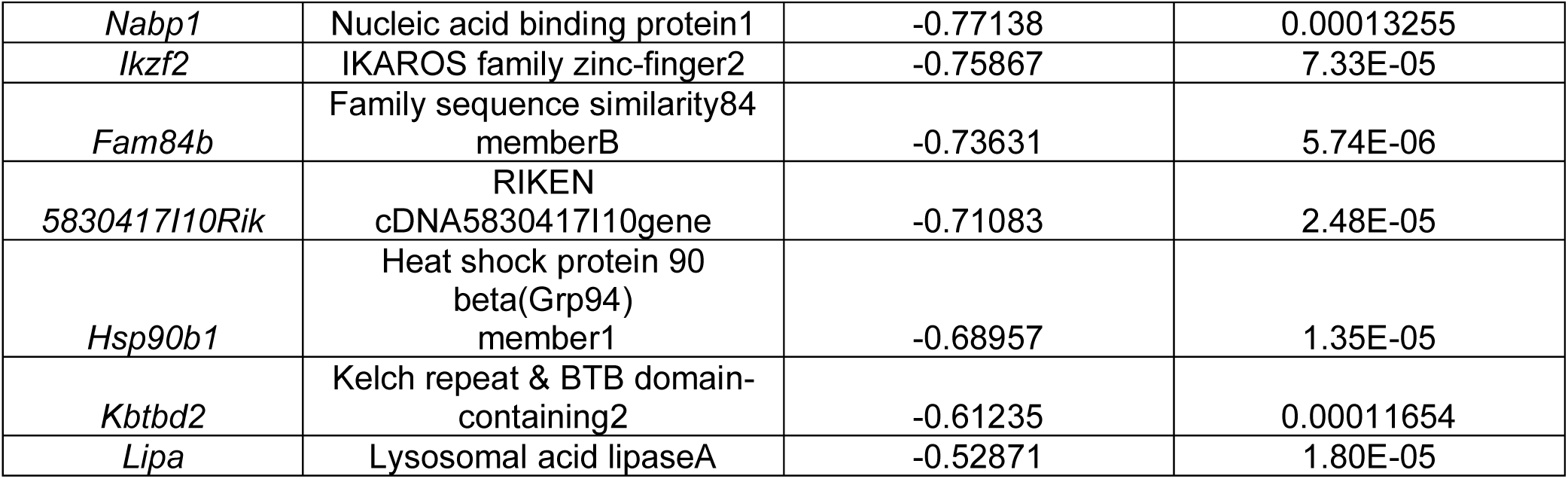
Differential Gene expression between WNV-infected KO mice compared to WNV-infected WT mice.

